# Human differentiated adipocytes can serve as surrogate mature adipocytes for adipocyte-derived extracellular vesicle analysis

**DOI:** 10.1101/2025.02.05.636729

**Authors:** Mangesh Dattu Hade, Bradley L. Butsch, Paola Loreto Palacio, Kim Truc Nguyen, Dharti Shantaram, Sabrena F. Noria, Stacy A. Brethauer, Bradley J. Needleman, Willa Hsueh, Eduardo Reátegui, Setty M. Magaña

## Abstract

Obesity is a growing global health concern, contributing to diseases such as cancer, autoimmune disorders, and neurodegenerative conditions. Adipose tissue dysfunction, characterized by abnormal adipokine secretion and chronic inflammation, plays a key role in these conditions. Adipose-derived extracellular vesicles (ADEVs) have emerged as critical mediators in obesity-related diseases. However, the study of mature adipocyte-derived EVs (mAdipo-EVs) is limited due to the short lifespan of mature adipocytes in culture, low EV yields, and the low abundance of these EV subpopulations in the circulation. Additionally, most studies rely on rodent models, which have differences in adipose tissue biology compared to humans. To overcome these challenges, we developed a standardized approach for differentiating human preadipocytes (preAdipos) into mature differentiated adipocytes (difAdipos), which produce high-yield, human adipocyte EVs (Adipo-EVs). Using visceral adipose tissue from bariatric surgical patients, we isolated the stromal vascular fraction (SVF) and differentiated preAdipos into difAdipos. Brightfield microscopy revealed that difAdipos exhibited morphological characteristics comparable to mature adipocytes (mAdipos) directly isolated from visceral adipose tissue, confirming their structural similarity. Additionally, qPCR analysis demonstrated decreased preadipocyte markers and increased mature adipocyte markers, further validating successful differentiation. Functionally, difAdipos exhibited lipolytic activity comparable to mAdipos, supporting their functional resemblance to native adipocytes. We then isolated preAdipo-EVs and difAdipo-EVs using tangential flow filtration and characterized them using bulk and single EV analysis. DifAdipo-EVs displayed classical EV and adipocyte-specific markers, with significant differences in biomarker expression compared to preAdipo-EVs. These findings demonstrate that difAdipos serve as a reliable surrogate for mature adipocytes, offering a consistent and scalable source of adipocyte-derived EVs for studying obesity and its associated disorders.

## Introduction

The widespread prevalence of obesity has escalated into a formidable global health emergency, placing significant burdens on public health systems worldwide ^1,2^. In 2019, an estimated 5 million obesity-related deaths occurred worldwide ^3^. Obesity is intricately linked to various metabolic disorders, including type 2 diabetes mellitus, cardiovascular diseases, and non-alcoholic fatty liver disease ^4–6^. Moreover, obesity is also implicated in the pathogenesis of several cancers, autoimmune diseases, and neurodegenerative disorders ^7,8^. The complex interplay between obesity and these diseases underscores the multifaceted role of adipose tissue in maintaining metabolic homeostasis and overall health ^9,10^. Adipose tissue, a dynamic endocrine organ, significantly influences energy balance, metabolic regulation, and immune responses ^11^. Dysfunctional adipose tissue, characterized by altered adipokine secretion, chronic inflammation, and impaired adipocyte differentiation, is a hallmark of obesity and a crucial factor in the emergence of obesity-related comorbidities ^12,13^. Recent research advances have illuminated the pivotal role of extracellular vesicles (EVs) in mediating intercellular communication and regulating metabolic processes ^14,15^. EVs are small, membrane-bound particles secreted by cells that transport bioactive molecules (e.g. proteins, lipids, and nucleic acids) to recipient cells, thereby altering recipient cell genotypic and phenotypic function ^14–16^.

Mounting evidence indicates that obesity is associated with enhanced production of adipose-derived EVs (ADEVs), which are instrumental in the pathogenesis of obesity and its related metabolic complications ^17–19^. ADEVs are implicated in various physiological and pathological processes, including inflammation, insulin resistance, and lipid metabolism ^19,20^.

Despite the growing interest in studying ADEVs, there is a dearth of human ADEV studies, partly due to the technical and biological limitations of obtaining EVs from mature human adipocytes (mAdipo-EVs). mAdipos exhibit short viability in culture, produce low yields of EVs, and are difficult to handle due to their nonadherent properties ^21–23^. To address these limitations, and with the end goal of developing a robust human translational paradigm for isolating mature adipocyte-derived EVs, we first developed a surrogate mAdipo model by differentiating human preAdipos into mature-like difAdipos with functional, genotypical, and phenotypical similarities to mature adipocytes. Secondly, we used the validated difApos as a reliable and surrogate source of human adipocyte EVs, demonstrating classic EV and adipose markers.

In summary, our study provides a reproducible and reliable source of human mature-like adipose EVs. By leveraging human-based models, we aim to provide a more relevant understanding of the role of adipocyte-derived EVs in obesity-related disorders, thus addressing the critical need for human-based obesity EV paradigms ^24^.

## Results

### Multiparametric isolation and characterization of ADEVs

We have developed a comprehensive pipeline for isolating and characterizing EVs from various human adipose tissue biospecimens^25^ (**Figure 1, Table S2)**. While most human EV biomarker studies have focused on biofluids or in vitro EV sources, isolating tissue derived EVs presents technical challenges that must be addressed to maintain the integrity of both the cells and the EVs^26^. To ensure our isolation method produced intact adipocyte derived EVs, we performed extensive bulk and single EV characterization, as detailed below in **Figure 1**.

**Figure 1.**
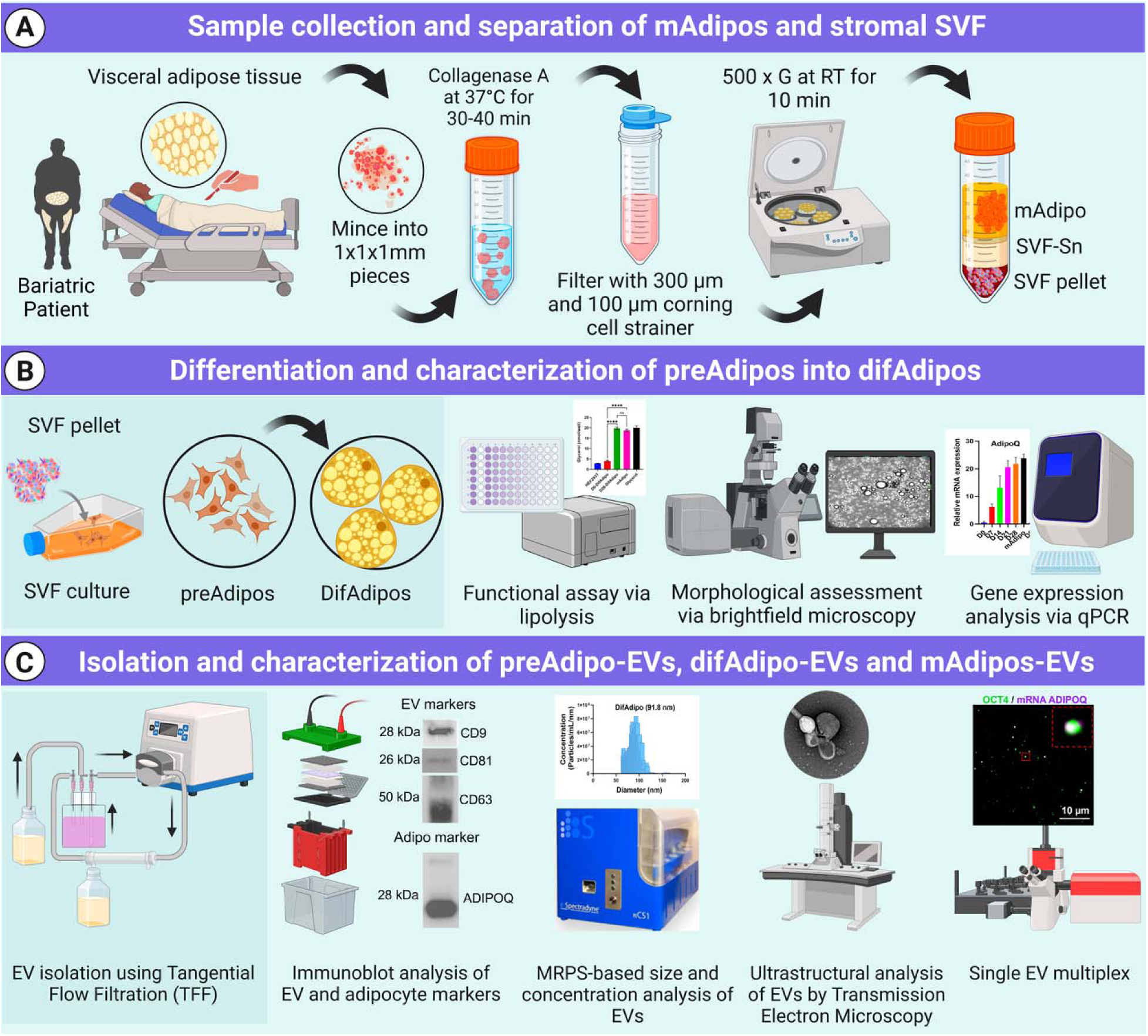
Workflow for the isolation, differentiation, and characterization of human preadipocytes, differentiated adipocytes, and their EVs. (A) Visceral adipose tissue obtained from bariatric patients was enzymatically digested using Collagenase A at 37 °C for 30–40 min, followed by filtration through 300 µm and 100 µm cell strainers. Subsequent centrifugation (500 × g, RT, 10 min) yielded three distinct fractions: stromal vascular fraction (SVF), SVF supernatant (SVF-SN), and mature adipocytes (mAdipos). The SVF-derived preadipocytes (preAdipos) were cultured and induced to differentiate into differentiated mature adipocytes (difAdipos) over 28 days. Conditioned media from adipocyte cultures were collected and processed using tangential flow filtration (TFF) for EV isolation. (B) Characterization of preAdipos, difAdipos and mAdipos included functional assays assessing lipolysis, morphological analysis via brightfield microscopy, and gene expression analysis using quantitative PCR. (C) Isolated EVs from preAdipos (preAdipo-EVs) and difAdipos (difAdipo-EVs) underwent extensive characterization, including immunoblot analysis of EV and adipocyte markers (CD9, CD81, CD63, and ADIPOQ), size distribution and concentration analysis via microfluidic resistive pulse sensing (MRPS), ultrastructural evaluation using transmission electron microscopy (TEM), and single-EV multiplexing analysis. extracellular vesicles (EVs); stromal vascular fraction (SVF), SVF supernatant (SVF-SN), mature adipocytes (mAdipos); preadipocytes (preAdipos); differentiated mature adipocytes (difAdipos); microfluidic resistive pulse sensing (MRPS); transmission electron microscopy (TEM)

### Visceral adipose-derived preadipocytes serve as a surrogate source for mature adipocytes

Our robust and systematic human-based protocol for isolating, differentiating, and characterizing SVF-derived preadipocytes yielded a reliable surrogate source of differentiated mature adipocytes (difAdipos) (see material and methods). Preadipocytes exhibited significant morphological changes during the differentiation process, accumulating larger lipid droplets and developing a mature phenotype, as confirmed by brightfield imaging **(Figure 2B)**.

**Figure 2.**
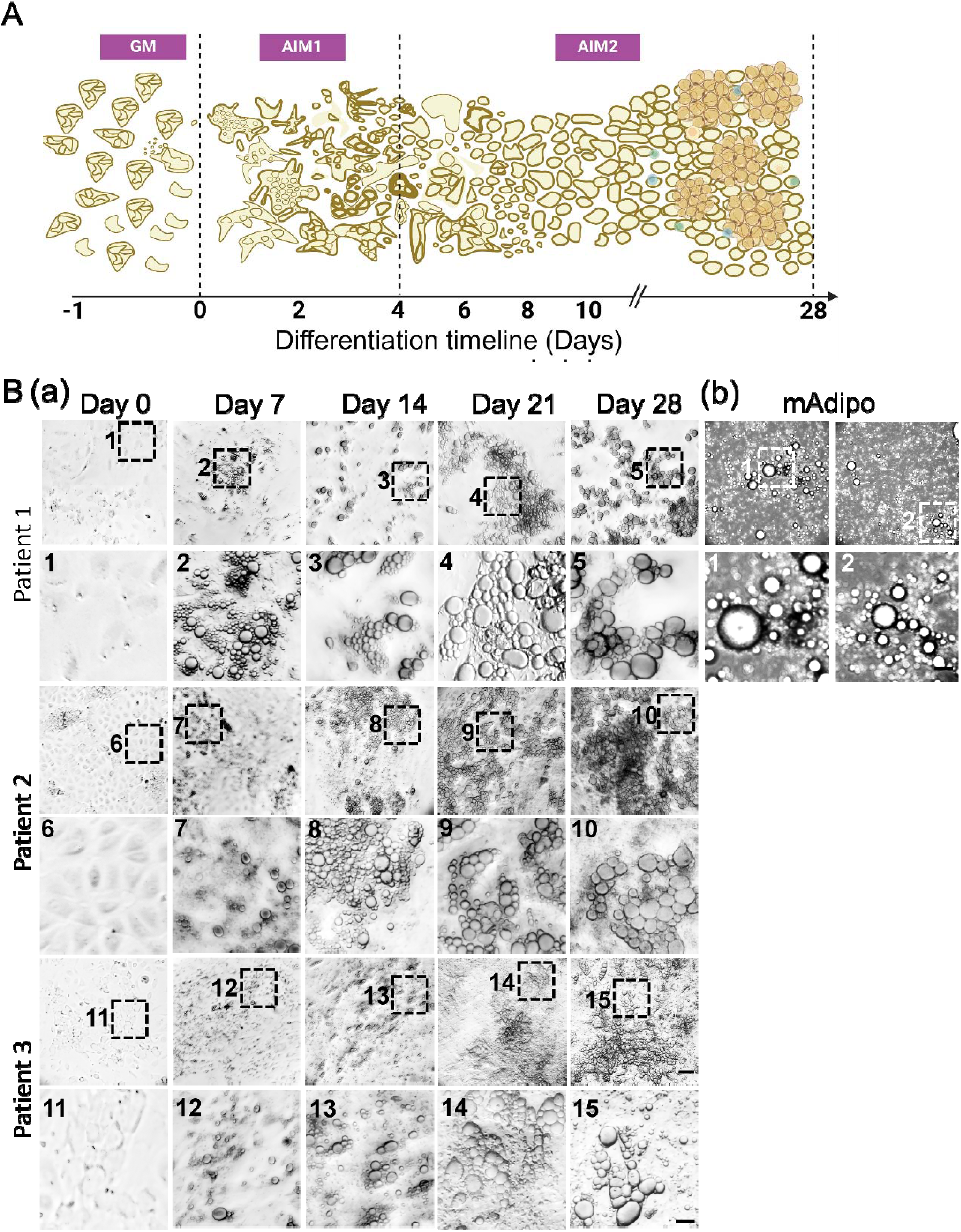
Human preadipocytes undergo differentiation and acquire a mature adipocyte phenotype. (A) Differentiation timeline: At Day -1, cells reached ∼80% confluency in growth medium (GM). Differentiation was initiated on Day 0 following a 24- hour incubation in GM by replacing the medium with adipogenic induction medium 1 (AIM1). The transition to adipogenic induction medium 2 (AIM2) was performed to facilitate further maturation, leading to the acquisition of a differentiated mature adipocyte (difAdipo) phenotype by Day 28. (B) Morphological changes during adipogenesis: (a) Representative brightfield microscopy images illustrate the progressive morphological changes occurring during preadipocyte differentiation into difAdipos. Lipid droplet accumulation becomes progressively evident throughout the differentiation process. (b) The morphology of difAdipos at Day 28 closely resembles that of primary mature adipocytes (mAdipos), which were directly isolated from patient-derived visceral adipose tissue and maintained in culture for 24 hr. Images are representative of three independent biological replicates (n = 3). Dashed boxes highlight selected regions, which are shown as enlarged high-magnification images in the subsequent panels to provide detailed visualization of cellular morphology. Scale bar = 50 µm for main images; 10 µm for enlarged high-magnification images. GM, growth medium; AIM1, adipogenic induction medium 1; AIM2, adipogenic induction medium 2. differentiated mature adipocyte (difAdipo); mature adipocytes (mAdipos)

### Differential expression of preadipocyte and mature adipocyte markers during adipocyte differentiation

We next investigated gene transcripts of known preAdipo markers (OCT4, PREF-1, and GATA3) ^27–30^ ^27^ and mature adipocyte markers (ADIPOQ, PLIN1, and PPARG) ^31^ at different time points during adipocyte differentiation (**Figure 3**). We hypothesized that difAdipos would acquire a genetic program similar to mAdipos and lose the undifferentiated transcript profile that characterizes preAdipos. As expected ^32,33^, OCT4 expression was low at Day 0 (D0), peaked at D7 (P< 0.001), and decreased as differentiation progressed and cells committed adipogenesis (**Figure 3A**).

**Figure 3.**
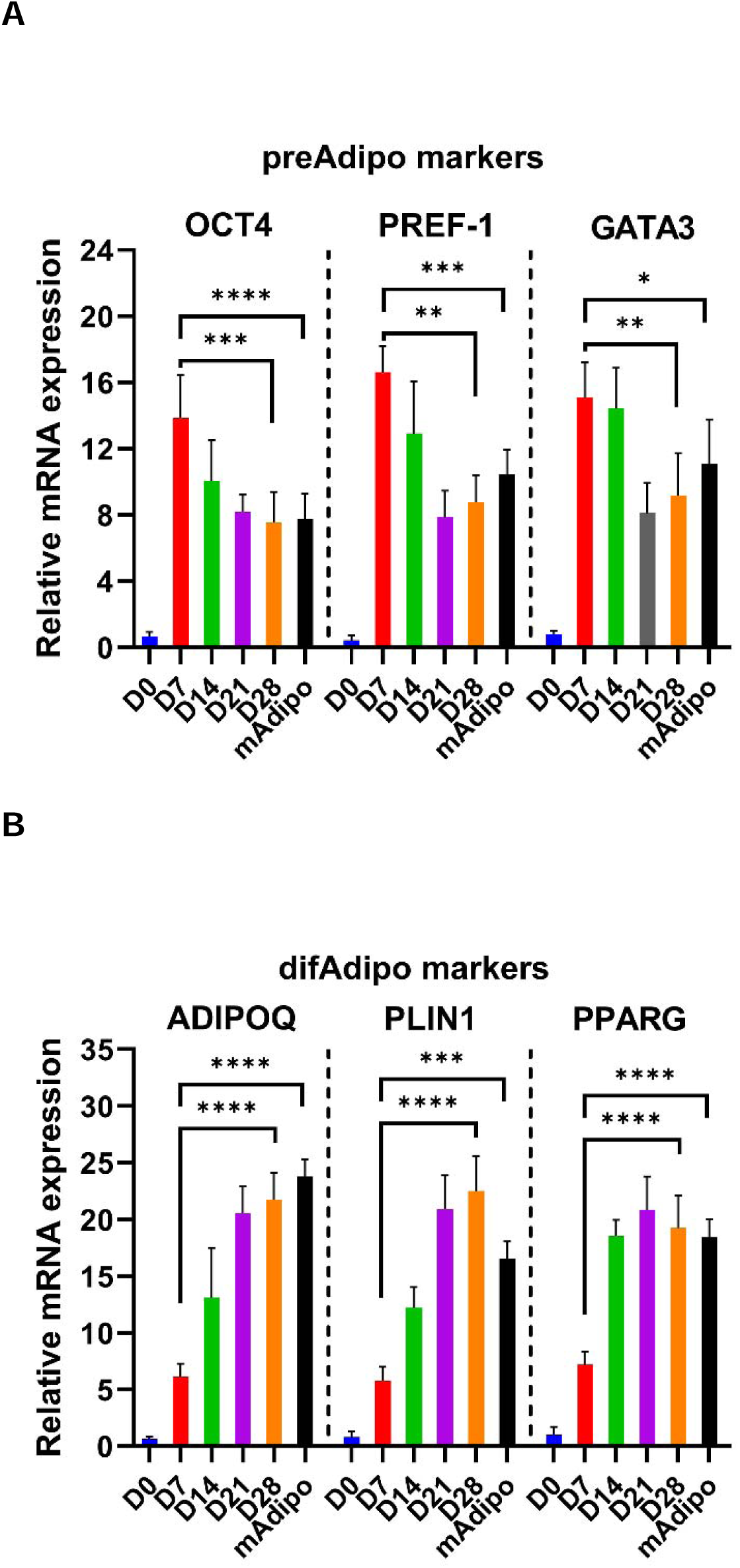
Transcript profile in preadipocytes, differentiated adipocytes and mature adipocyte. **(A)** Relative mRNA expression levels of preadipocyte markers (OCT4, PREF1, GATA3) and **(B)** mature adipocyte markers (ADIPOQ, PLIN1, PPARG) were measured at Day 0 (D0), D7, D14, D21, D28, and in terminally differentiated mAdipo. OCT4 expression peaked at D7 and decreased thereafter, indicating early differentiation. PREF-1 expression increased to a peak at D7 and decreased towards D28, consistent with its inhibitory role in adipogenesis. GATA3 expression peaked at D7 and D14 and remained elevated in mAdipo, indicating its involvement in early differentiation. Adipocyte markers ADIPOQ, PLIN1, and PPARG exhibited a progressive increase, peaking at D28 and in mAdipo, reflecting mature adipocyte function (n=3, biological replicated). The statistical analysis was performed using ordinary one-way ANOVA with Bartlett’s test. OCT4, octamer-binding transcription factor 4; PREF-1, preadipocyte factor 1; GATA3, GATA-binding protein 3; ADIPOQ, adiponectin; PLIN, perilipin; PPARG, peroxisome proliferator-activated receptor gamma. A value of P < 0.05 was deemed statistically significant (*< 0.05; **< 0.01; ***< 0.001; ****< 0.0001).

We saw a similar pattern in PREF-1 expression levels. PREF-1 is an inhibitor of adipogenesis and it is downregulated as cells undergo differentiation ^33^. Consistent with this observation, preAdipo PREF-1 levels were relatively low at D0, peaked at D7(P< 0.0001), and were decreased by D28 and in mAdipos (**Figure 3A**). In contrast to OCT4 and PREF-1, GATA3 levels persisted in both the early and intermediate differentiation stages. GATA3 expression peaked at D7 (P< 0.01) and D14, decreased through D21 and D28, but remained elevated in mAdipo compared to D28 (**Figure 3A**). This expression pattern is consistent with studies showing GATA3’s involvement in regulating adipogenesis ^34,35^.

Concurrently, the expression of the mature adipocyte markers ADIPOQ, PLIN1, and PPARG, showed a clear increasing trend during differentiation, as previously reported ^35–37^. ADIPOQ expression was low at D0 and D7, increased steadily from D14 onwards, and peaked at D28 (P< 0.0001), with levels comparable to those seen in our mAdipos (**Figure 3B**). PLIN1 plays a crucial role in regulating lipid storage and mobilization by protecting lipid droplets from lipolysis in adipocytes ^38^. We observed gradually increasing levels of PLIN1 expression from D0 onward, significantly increasing at D21 and D28 (P< 0.0001), signifying lipid accumulation and adipocyte maturation.

Similarly, PPARG expression progressively increased from D0 to D28, with the highest expression observed at D21 and D28 (P< 0.0001), and slightly lower expression in mAdipo, highlighting its crucial role as a regulator of adipogenesis (**Figure 3B**). In summary, the dynamic transcriptional program we observed reflects the successful differentiation of preadipocytes into difAdipos exhibiting mature adipocyte transcriptional profiles.

### Differentiated adipocytes exhibit lipolytic function comparable to mature adipocytes

Preadipocytes differentiated in vitro may not retain the same biophysical properties of mature adipocytes that are differentiated in vivo^39^. Therefore, we next sought to functionally validate the mAdipo phenotype of our difAdipos by investigating their ability to perform the prototypic mature adipocyte function of lipolysis ^40–42^. Cells were stimulated with the lipolytic agent, isoproterenol, and lipolysis was assessed by quantifying the stimulated release of glycerol into the culture medium (**Figure 4**). We hypothesized that the lipolytic activity of difAdipos would not differ from that of mAdipos, suggesting that our difAdipos could serve as functional surrogates of mAdipos.

**Figure 4.**
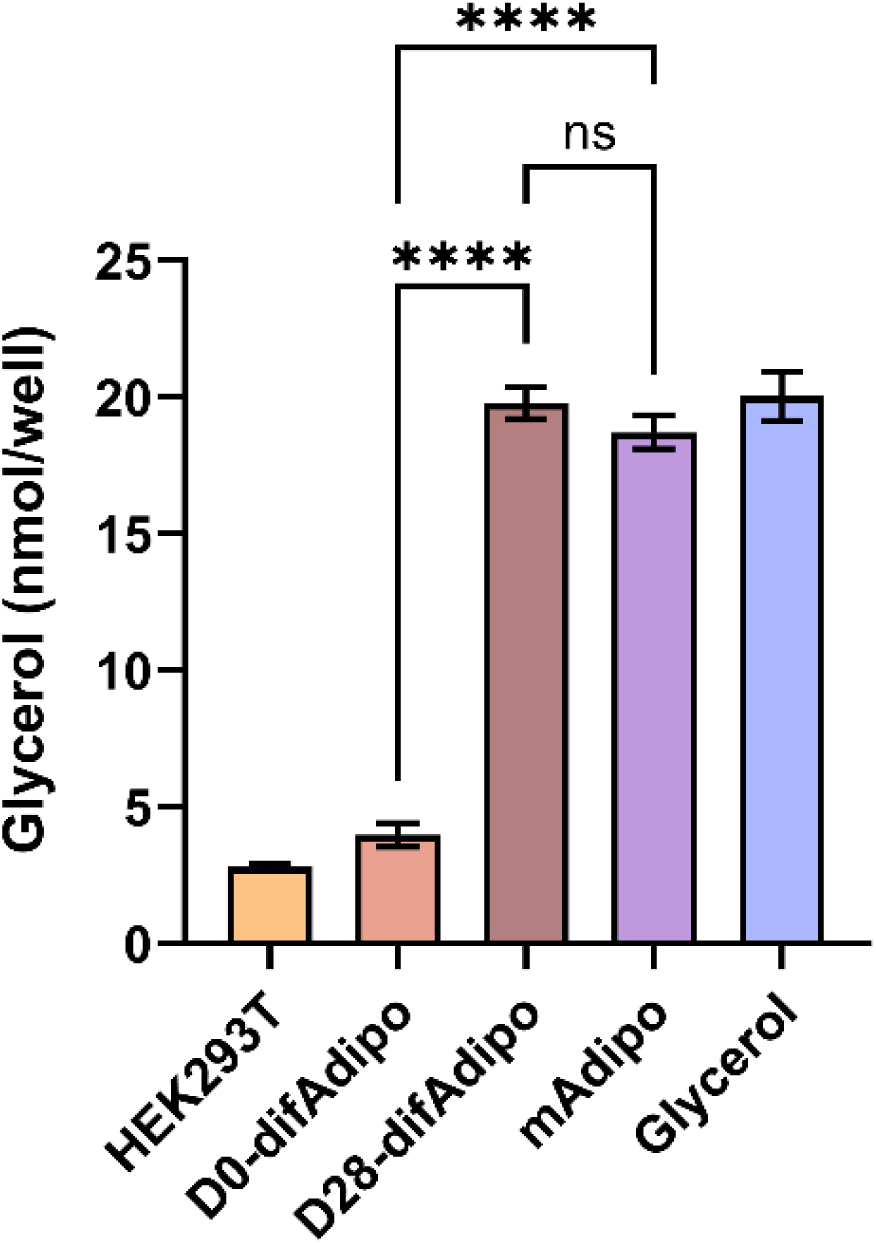
Differentiated adipocytes (difAdipos) exhibit lipolytic function comparable to mature adipocytes (mAdipos). Lipolytic activity was assessed in preAdipos, difAdipos, and mAdipos by stimulating cells with isoproterenol and measuring glycerol release. D0 difAdipos reflect undifferentiated adipocytes and exhibited low-level lipolytic activity similar to HEK293T cells (negative control). D28 difAdipos exhibited lipolytic activity comparable to mAdipos and glycerol (positive control), validating the differentiation protocol and confirming that difAdipos functionally resemble mature adipocytes in their lipolytic activity. (n=3, biological replicated) Statistical analysis was performed using one-way ANOVA with Sidaks multiple comparison test. A value of P < 0.05 was deemed statistically significant (*< 0.05; **< 0.01; ***< 0.001; ****< 0.0001).

Compared to undifferentiated D0 preadipocytes, D28 difAdipos achieved a comparable degree of lipolytic activity as mAdipos. These findings demonstrate our ability to generate human difAdipos that are not only morphologically and genotypically comparable to mAdipos, but also functionally resemble mAdipos.

### Multiparametric characterization of EVs from differentiated adipocytes

After validating our difAdipos as robust surrogates for mAdipos, we next sought to characterize EVs from matched preAdipos, difAdipos, and mAdipos obtained from the same individual (**Figure 5A**). Microfluidic resistive pulse sensing analysis revealed a relatively narrow particle size distribution among the EVs from preAdipo, difAdipo, and mAdipo. Median particle diameters were similar across the three Adipo populations (88.5 nm for preAdipo, 91.8 nm for difAdipo, and 85.1 nm for mAdipo). All three ADEV sources had similar mean particle concentrations: 6.26E+10 particles/mL for mAdipos, 3.47E+10 particles/mL for preAdipos and 2.33E+10 particles/mL for difAdipos **(Figure 5A).** These results suggest that our differentiation protocol did not alter the ability of difAdipos to generate small EVs that were similar in size and concentration to native populations (ie. preAdipos and mAdipos), which did not undergo in vitro maturation.

**Figure 5.**
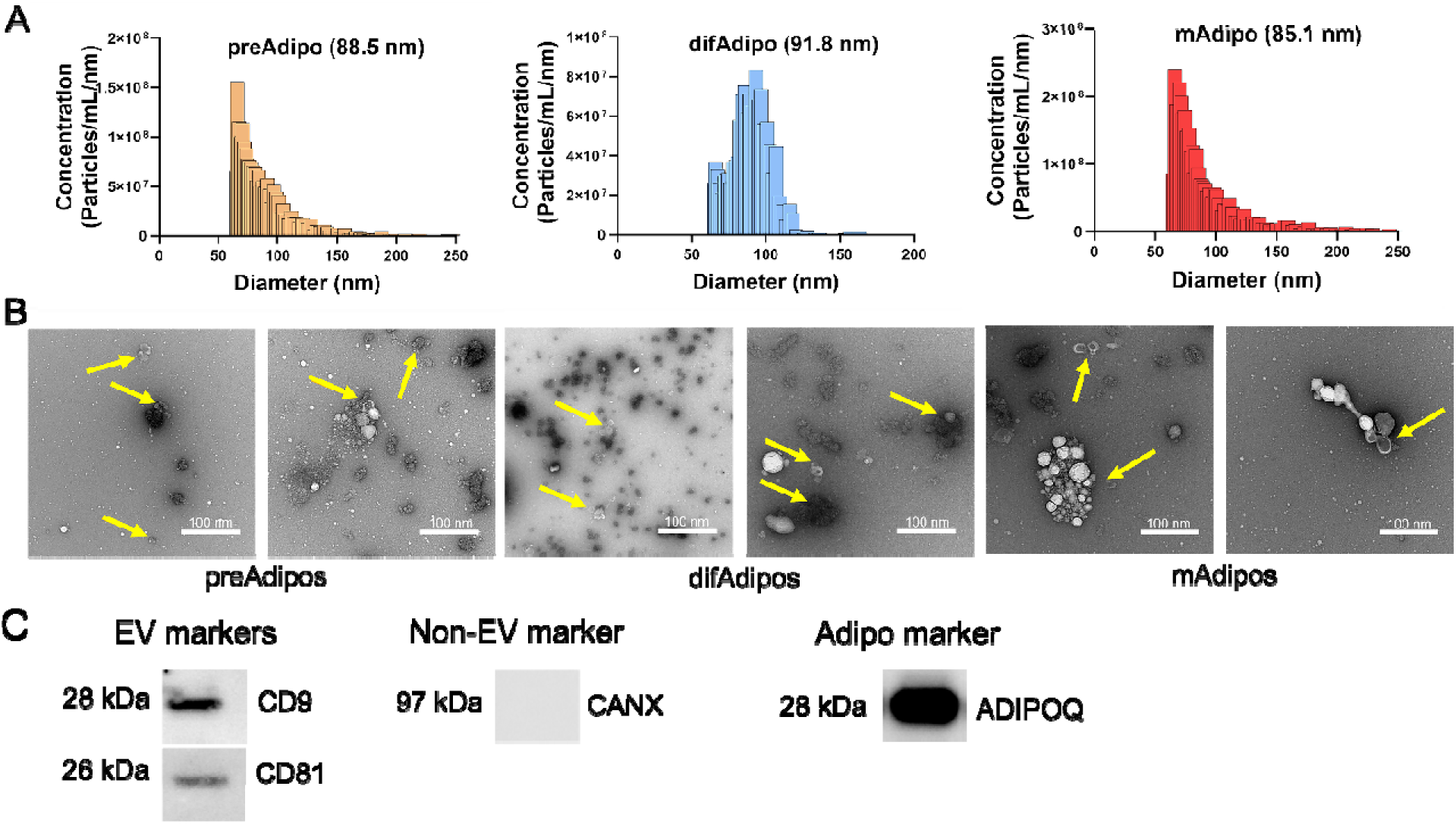
Multiparametric characterization of matched adipose tissue-derived EVs from the same bariatric patients. (A) MRPS analysis of EVs from preAdipo, difAdipo, and mAdipo from the same individuals (n = 3 biological replicated). Median particle diameters: 88.5 nm (preAdipo), 91.8 nm (difAdipo), 85.1 nm (mAdipo). Mean particle concentrations: 3.47E+10 particles/mL (preAdipo), 2.33E+10 particles/mL (difAdipo), 6.26E+10 particles/mL (mAdipo). (B) TEM imaging showed a polydispersed population of particles, including small EVs with a characteristic ‘cup-shaped’ morphology. (C) Immunoblot analysis of EVs from difAdipo reveals heterogeneous expression of typical EV (CD9, CD81), non-EV (CANX), and adipose-specific (ADIPOQ) markers. Full-size uncropped blot images for panel (C) are provided in Supplementary Figure S1. MRPS, microfluidic resistive pulse sensing; TEM, transmission electron microscopy.

TEM revealed a polydispersed and prominent population of small EVs with the characteristic artefactual ‘cup-shaped’ morphology **(Figure 5B).** Immunoblotting of difAdipo-EVs demonstrated a heterogeneous expression of the typical EV markers CD9, CD81, and the absence of the non-EV marker calnexin (CANX) **(Figure 5C, S1)**. The adipocyte marker, ADIPOQ, was also detected (**Figure 5C, S1)**. Collectively, these findings highlight the diversity of tissue-derived EV populations from varied adipocyte sources within the same individual and at different stages of adipocyte differentiation.

### High-resolution, simultaneous detection of human adipose-derived EV proteins and RNA using total internal reflection fluorescence (TIRF) microscopy

TIRF microscopy was utilized to achieve simultaneous detection of human adipose-derived EV proteins and RNA and colocalization of the same (**Figure 6 A, B).** First, we investigated the colocalization of EV surface proteins and EV mRNA cargo in preAdipo-EVs and difAdipo-EVs to confirm the specificity of adipocyte-EVs. DifAdipo-EVs revealed high colocalization between OCT4 protein and ADIPOQ mRNA, confirming the adipocyte-specificity of the EVs. The tetraspanin marker CD63 showed less to no colocalization with other biomarkers (Figure 6A), underscoring the heterogeneous expression profile of tetraspanins. Similar results were observed for preAdipo-EVs (results not shown).

**Figure 6.**
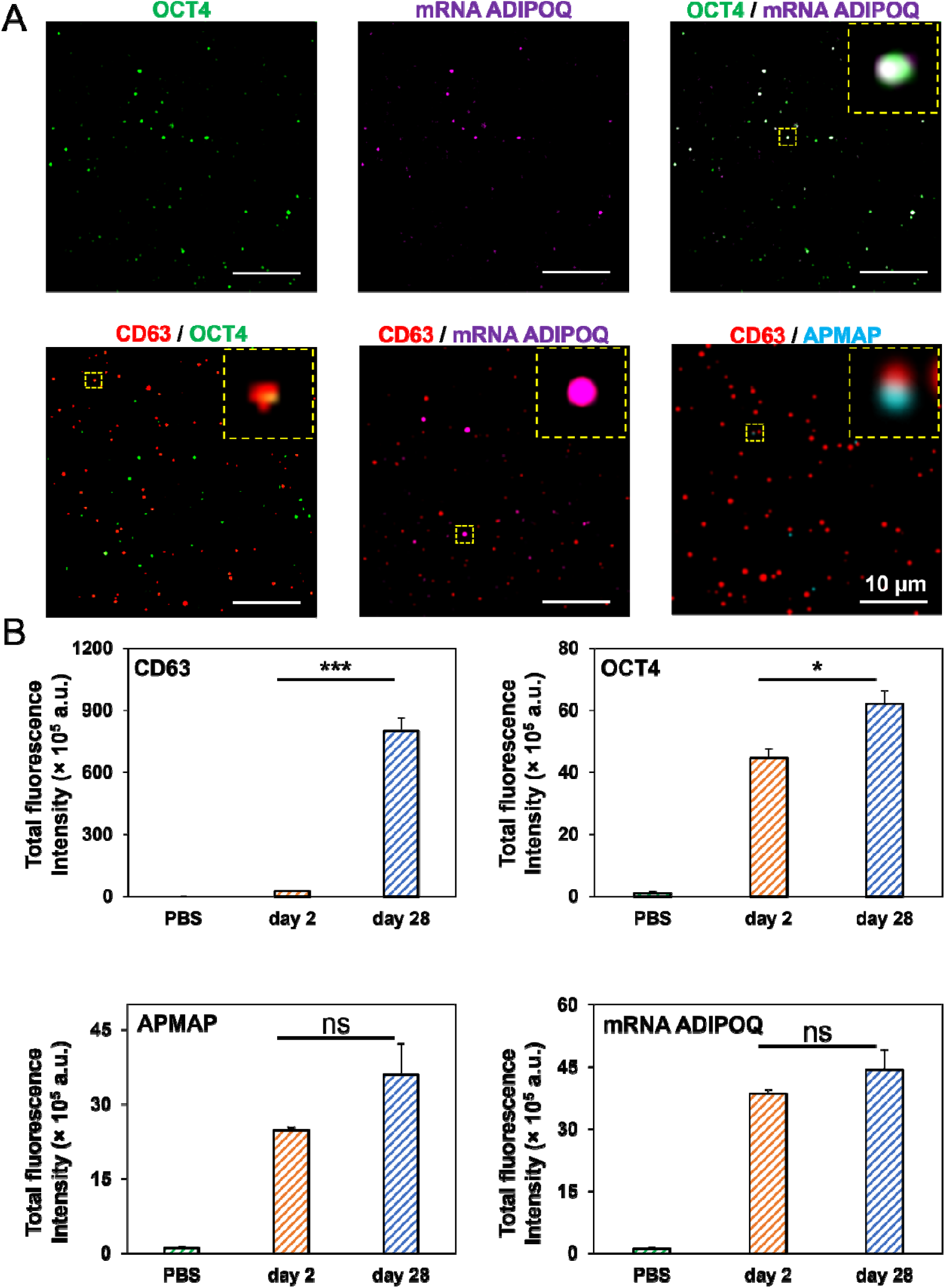
Single EV multiparametric characterization comparing preAdipo-EVs (day 2) and difAdipo-EVs (day 28). (A) Representative TIRFM images illustrating difAdio-EVs were captured by anti-CD9 and anti-CD63 antibodies and immobilized on a biochip surface. Detection antibodies against CD63, OCT4, and APMAP were used to identify EV surface proteins. A fluorescence-labeled molecular beacon detected the mRNA ADIPOQ within those EVs. The merged image reveals difAdipo-EVs defined by colocalization of OCT4 and mRNA ADIPOQ, CD63 and OCT4, CD63 and mRNA ADIPOQ, CD63 and APMAP Scale 10 µm (B) The relative fluorescence intensity of EVs collected at day 2 and day 28 was analyzed for the tetraspanin marker CD63, the undifferentiated cell marker OCT4, the adipocyte plasma membrane-associated protein APMAP, and the adipose tissue gene marker mRNA ADIPOQ. OCT4, octamer-binding transcription factor 4; ADIPOQ, adiponectin; APMAP, astrocyte membrane-associated protein. Statistical analysis was performed using students test. A value of P < 0.05 was deemed statistically significant (*< 0.05; **< 0.01; ***< 0.001; ****< 0.0001).

Next, we quantified the relative fluorescence intensity of EV surface markers and mRNA cargo (**Figure 6B**). DifAdipo-EVs collected on day 28, compared to preAdipo-EVs collected on day 2, expressed a significantly higher level of the canonical EV marker CD63 (***< 0.001). Interestingly, the protein marker for undifferentiated adipocytes, OCT4, was also significantly higher on difAdipo-EVs than preAdipo-EVs (*< 0.05). The expression levels of adipose tissue-specific protein and gene markers, such as adipocyte plasma membrane-associated protein (APMAP) and ADIPOQ, were similar between the difAdipo-EVs and preAdipo-EVs. In summary, APMAP and ADIPOQ exhibited stable expression levels across the maturation process, while CD63 and OCT4 were dynamically regulated during the same differentiation process.

## Discussion

Adipose-derived EVs are increasingly recognized as critical orchestrators in obesity-associated diseases^43^. Limited studies have characterized human adipocyte EVs^44–48^ , however, no study has systematically studied paired difAdipo-EVs and mAdipo-EVs at bulk and single EV resolutions. Herein, we present a reproducible and scalable surrogate source of human mature-like adipocytes (difAdipos) that exhibit phenotypic, genotypic, and functional properties paralleling those of paired mature adipocytes (mAdipos). Furthermore, we demonstrate that in vitro difAdipos can release ADEVs comparable to ADEVs released by mature adipocytes. Finally, we provide the first high-resolution, single EV analysis of human difAdipo-EVs, demonstrating simultaneous adipocyte-enriched gene transcript and surface protein markers.

Due to recognized cross-species differences^49,50^ between mouse and human adipose biology, adipocyte EVs from rodent models have limited direct human translational utility. Several alternate approaches to circumvent the cross-species differences include studying in vitro differentiated human adipocytes, adipose tissue explants, and circulating adipocyte-enriched EV subpopulations in human biofluids^51^. Kranendonk and colleagues were amongst the earliest groups to characterize human adipocyte EVs (defined as adiponectin and fatty-acid binding protein 4 positive EVs), isolated from the Human Simpson Golabi Behmel Syndrome (SGBS) cell line^52^ . More recently, a study by Clement and collaborators isolated primary human adipocytes from dermolipectomy-derived subcutaneous adipose tissue and demonstrated the functional ability of isolated human adipocyte EVs to metabolically remodel melanoma cells into a more invasive phenotype^53^. Several human ADEV studies have utilized tissue explants for ADEVs rather than mature adipocytes^48,54^ . Human adipose tissue explants have the advantage of overcoming the cross-species limitations of mouse models but confound the ability to investigate the potential pathobiological role of EVs shed solely from mature adipocytes^55^which are the most abundant and primary functional metabolic cells within visceral adipose tissue^46,56^.

A less invasive and more feasible approach for studying human ADEVs involves interrogating the circulating pool of all EVs and attempting to enrich for EV subpopulations that contain ‘adipocyte-specific’ markers. Connolly and colleagues were the first to confirm human EVs containing adipose-enriched markers within the circulating pool of plasma-derived EVs from obese individuals^57^ . In this study, the authors applied contemporaneous ISEV guidelines^58^ for EV research and used complementary EV isolation and enrichment methods to attempt to account for non-vesicular co-isolates and non-adipose EV populations.

One of the formidable challenges in studying cell and tissue-specific EVs, such as ADEVs, arises from their low abundance and high signal-to-noise ratio^55,59^ . Adipose EVs are estimated to comprise >80% of tissue-derived EVs in human plasma and can be the primary source of circulating miRNAs; however, tissue-derived EVs comprise <0.2 % of the circulating human plasma pool ^60–62^. This limitation underscores the critical need for more advanced methods to study low-abundant, rare EV subpopulations. To address this limitation, we first produced a surrogate source of functional mature adipocytes in vitro and demonstrated their ability to shed EVs comparable in adipocyte-specific markers to EVs shed by adipocytes that matured in vivo.

To our knowledge, no studies have characterized the biophysical properties of human adipose-derived EVs in the pre- and post-adipogenic phases from paired human biospecimens. One study conducted a systematic phenotypic comparison between immature and mature mouse ADEVs^63^. In this study, the authors characterized the biophysical properties of pre- and post-adipogenic adipocyte-derived EVs from 3T3-L1 cells. Similar to the Connolly study, we found the highest expression of the preAdipo marker PREF-1 in the earliest stages of differentiation and the highest expression of the mAdipo marker ADIPOQ in the later stages of maturation. They also found increased secretion of small EVs in the early stages of differentiation, slightly increased CD9 and CD63 protein via time-resolved fluorescence, and a higher proportion of phospholipids associated with cell signaling. Interestingly, the authors found EV-unique characteristics that differed from the 3T3-L1 parental cell, suggesting modifications that could impact EV-mediated intercellular communication^63^.

Finally, we applied single EV multiplexing analytics to validate the adipocyte-specificity of our difAdipo-EVs. Colocalization of EV surface proteins and internal cargo can increase sensitivity for detecting low-abundance EV subpopulations and can also enhance the biomarker utility of EVs to detect earlier stages of disease^64,65^ . Therefore, our single ADEV multiplexing platform holds great promise for investigating novel and robust, adipocyte-specific EVs in obesity-associated diseases.

## Conclusion

Our study presents a novel paradigm for advancing the translational application of human ADEVs by using human difAdipos as a surrogate for mature adipocytes, thereby augmenting the harvesting of viable human-derived, mature adipocyte EVs. The model established provides a high-yield, functionally equivalent source of EVs, facilitating more consistent and relevant research into the role of ADEVs in obesity and metabolic diseases. Our findings underscore the importance of adipocyte differentiation on EV content and function, revealing significant differences in biomarker profiles between preAdipo-EVs and difAdipo-EVs. This differentiation may impact the bioactive cargo within the EVs, influencing recipient cell behavior and contributing to metabolic complications associated with obesity.

Future research should explore the specific mechanisms by which difAdipo-EVs influence metabolic processes and their potential as biomarkers or therapeutic targets in obesity-related conditions. This study herein lays the groundwork for developing therapeutic strategies that modulate ADEVs’ effects, offering promising avenues for mitigating the adverse consequences of obesity on metabolic health and reducing the risk of associated diseases.

## Materials and methods

### Study participants

This study complies with all relevant ethical guidelines. The Ohio State University Institutional Review Board (IRB# 2014H0471) approved the study, and all participants provided written informed consent. Visceral adipose tissue (VAT) samples were collected from patients (n = 11, mean age: 41.6 years, SD: 12.98; mean BMI: 40.97, SD: 6.00) undergoing elective bariatric surgery at The Ohio State University (OSU) Center for Minimally Invasive Surgery. Clinical data are summarized in Table S1.

### Adipose tissue dissociation and mature adipocyte culture for EV isolation

Adipose tissue samples were collected in the operating room during the bariatric surgery and processed within 1 hr using collagenase digestion, as previously described^25^. The cell suspension was centrifuged at 500 x g for 10 min to generate three distinct phases (mature adipocytes, the stromal vascular fraction (SVF), and the SVF secretome). Floating mature adipocytes were carefully removed from the top layer and kept for culture. The supernatant was then carefully aspirated to avoid disturbing the SVF pellet, which was resuspended in sorting media for the pre-adipocyte isolation and differentiation protocol (see below). Mature adipocytes were cultured for 24 hr in Dulbecco’s Modified Eagle Medium/Ham’s Nutrient Mixture F-12 (DMEM/F12), supplemented with 40% M199 basal medium with Earle’s salts and L-glutamine (MCDB201), 2% fetal bovine serum (FBS), 1x penicillin/streptomycin, 1 nM dexamethasone, 0.1 mM L-ascorbic acid 2-phosphate (LAAP), 1x Insulin-Transferrin-Selenium (ITS) Mix, 1x linoleic acid-albumin, 5 μg/μL insulin, and 1 nM triiodothyronine (T3).

### Preadipocyte (preAdipo) isolation and differentiation into differentiated mature adipocytes (difAdipos)

The resuspended SVF pellet was filtered through a 100 µm cell strainer into fresh tubes, repeating the washing step to ensure thorough isolation of SVF cells. Following another round of centrifugation (at 1200 rpm for 10 min at RT or 4 °C), and aspiration of the supernatant, the SVF pellet was treated with ACK lysis buffer on ice for three minutes to lyse any remaining red blood cells, followed by dilution and filtration through a 40 µm cell strainer. Finally, the SVF pellet containing cells were re-suspended in sorting media and seeded into culture plates until the cells reached approximately 80% confluency.

The differentiation process commenced with a 24 hr incubation in growth media, designated as day 0. Throughout the differentiation period, media was replaced every 48 hr. Cells were maintained for up to 28 days in adipogenic induction media 1 and 2 (AIM1 and AIM 2). For media composition, see Table S3.

### ADEV isolation via tangential flow filtration (TFF)

Culture media from PreAdipo, difAdipo, and mAdipo were first filtered through 0.2 μm filters and concentrated to a final volume of 5 mL. The concentrated media underwent diafiltration using tangential flow filtration (TFF) with a 500 kDa filter at a flow rate of 35 mL/min, as described previously^25^. After TFF, the retentates were further concentrated to 100 μL using 30 kDa molecular weight cutoff (MWCO) centrifugal filter units (MilliporeSigma Amicon Ultra, Fisher Scientific) by centrifugation at 4000 × g for 30 minutes.

### Microfluidic resistive pulse sensing (MRPS)

The concentration and size distribution of particles were assessed using microfluidic resistive pulse sensing (MRPS) on the Spectradyne nCS1 instrument (Spectradyne, Torrance, CA). Measurements were conducted with C-400 polydimethylsiloxane cartridges, enabling detection within an approximate size range of 65–400 nm^25^. Data processing and analysis were performed using the nCS1 Data Analyzer software (Spectradyne, Torrance, CA).

### Western blotting

EV samples were lysed using radioimmunoprecipitation assay (RIPA) buffer (Thermo Scientific) supplemented with protease and phosphatase inhibitors (Thermo Scientific) and incubated on ice for 15 min. Protein concentration was measured with the Micro BCA™ Protein Assay Kit (Thermo Scientific). Equal protein amounts were mixed with Laemmli buffer containing 2-mercaptoethanol (Sigma-Aldrich) and separated via SDS-PAGE on 4–20% Mini-PROTEAN® TGX Stain-Free gels (Bio-Rad). The proteins were then transferred onto polyvinylidene fluoride (PVDF) membranes (Bio-Rad). After blocking, membranes were incubated overnight at 4 °C with primary antibodies prepared in TBS-T, followed by a 1 hr incubation with horseradish peroxidase-conjugated secondary antibodies at room temperature. Protein detection was carried out using enhanced chemiluminescence (ECL) reagents (Bio-Rad), and signals were visualized using the Bio-Rad ChemiDoc™ MP imaging system.

### Quantification of real-time polymerase chain reaction (qRT-PCR)

Total RNA was extracted using the RNeasy Mini Kit (Qiagen) following the manufacturer’s protocol and quantified with a NanoDrop spectrophotometer. The purified RNA was then reverse-transcribed into complementary DNA (cDNA) using the High-Capacity cDNA Reverse Transcription Kit (Applied Biosystems). Quantitative real-time PCR was conducted using PowerUp™ SYBR™ Green Master Mix (Thermo Fisher) on an Applied Biosystems real-time PCR system. The forward and reverse primer sequences are provided in **Table S4**. β-actin served as the internal reference gene for normalization.

### Lipolysis assay

To investigate mature adipocyte functional activity, a lipolysis assay was performed using the Lipolysis Assay Kit (Abcam) per the manufacturer’s instructions. Day 0 (preAdipo), day 28 (difAdipo), and cultured HEK293 (negative control) cells were washed twice with Lipolysis Wash Buffer, then stimulated with 100 nM isoproterenol for 4 hr. Absorbance at 570 nm was measured using the SpectraMax iD3 (Molecular Devices), and glycerol concentration was quantified using a standard curve.

### Transmission electron microscopy (TEM)

TEM was performed as previously described^66^. Briefly, TEM grids were first treated with plasma for high surface EVs absorption. Subsequently, 10 µL of EVs samples were drop-casted onto the plasma-treated surface. The samples were incubated for 1 minute and then blotted with filter paper to remove excess liquid. The TEM grids were washed twice in DI water and stained with UranyLess EM contrast stain (Electron Microscopy Science) for 22 sec. The TEM grids were dried overnight prior to imaging. TEM images were obtained using Tecnai TF-20 microscope under 200kV, bright-field imaging mode.

### Single EV analysis using total internal reflection fluorescence microscopy (TIRF)

The isolated EVs were captured and analyzed on a gold functionalized biochip following our previous protocol ^25^. Briefly, the EVs collected on day 2 (preAdipo-EVs) and day 28 (difAdipo-EVs) were permeabilized and hybridized with molecular beacons (**Table S5**) in 0.5 × Tris EDTA (TE) buffer at 0.2 µM concentration for 2 hr at 37 °C in a dark environment to facilitate molecular beacon hybridization to the target RNA. The EV samples were incubated on the biochip surface and captured by functionalized anti-CD63 and anti-CD9 antibodies. Proteins were detected using fluorescent-dye-conjugated antibodies. Finally, a 10 × 10 array of images was acquired via total internal reflection fluorescence microscope (TIRFM; Nikon, Melville, NY) for each well. Relative and total fluorescence intensities of the sample were obtained from custom-built algorithms that were previously reported ^67,68^

### Data availability

All data generated or analyzed during this study are included in this manuscript and its supplementary information file. Further inquiries can be directed to the corresponding author.

## Author contributions

Conceptualization: M.D.H., BLB., P.L.P., W.H., E.R., S.M.M,

Study design and funding acquisition: E.R., S.M.M.

Investigation and performed research: M.D.H., B.L.B., P.L.P., D.S., K.T.N., S.N., S.A.B., B.J.N.

Organization and data analysis: M.D.H., B.L.B, K.T.N

Completed the original draft of the manuscript: M.D.H., B.L.B., S.M.M

Data interpretation, revision, and manuscript correction: M.D.H., B.L.B., K.T.N., E.R., S.M.M.

All authors approved the final version of the manuscript.

## Additional information

Correspondence and requests for materials should be addressed to Senior Corresponding author.

## Acknowledgments

SMM was supported by intramural funding from the Abigail Wexner Research Institute at Nationwide Children’s Hospital and NINDS (5K12NS098482-05). ER was supported by National Institutes of Health (NIH) grants UG3/UH3TR002884 and U18TR003807. The content is solely the responsibility of the authors and does not necessarily represent the official views of the National Institutes of Health.

