## Supplementary information for "Human differentiated adipocytes can serve as surrogate mature adipocytes for adipocyte-derived extracellular vesicle analysis"

**Running Title:** Human differentiated adipocyte extracellular vesicles

***Corresponding Author:** Setty Magaña, MD, PhD

Address:	Nationwide Children’s Hospital

Department of Pediatrics, Division of Neurology

700 Children’s Drive

Columbus, OH 43205

**Table S1:** **Participants demographics.**

| **ID** | **Age** | **Sex** | **Ethnicity** | **BMI (kg/m²)** |
| --- | --- | --- | --- | --- |
| **1** | 31 | F | African American | 49.51 |
| **2** | 21 | F | Caucasian | 42.64 |
| **3** | 42 | F | Caucasian | 38.42 |
| **4** | 32 | F | Caucasian | 40.22 |
| **5** | 31 | F | Caucasian | 47.12 |
| **6** | 43 | F | Caucasian | 34.93 |
| **7** | 60 | F | Caucasian | 38.58 |
| **8** | 35 | F | African American | 46.83 |
| **9** | 43 | F | Caucasian | 34.67 |
| **10** | 55 | F | Caucasian | 30.18 |
| **11** | 65 | F | Caucasian | 47.59 |

**Table S2. Definition and source of adipose cells used in this study.**

| Abbreviation | Definition | Study Source | EV population |
| --- | --- | --- | --- |
| preAdipos | preadipocytes | Derived from the stromal vascular fraction   - Differentiated in vitro for 28 days under adipogenic induction and growth media | preAdipo-EVs |
| difAdipos | In vitro differentiated adipocytes (surrogate source for mature/native adipocytes) | preAdipos | difAdipo-EVs |
| mAdipos | In vivo (native) differentiated mature adipocytes | Derived from adipose tissue dissociation   - Cultured ex vivo for 24 hours | mAdipo-EVs |

**Table S3: Media Composition**

| **AIM1** | **Ingredients** | **Stock Concentration** | **Final Concentration** |
| --- | --- | --- | --- |
|  | DMEM/low glucose |  | 60% |
|  | MCDB201 Media |  | 40% |
|  | FBS |  | 2% |
|  | Penicillin/Streptomycin (P/S) | 100x | 1x |
|  | Dexamethasone (DXM) | 10uM | 1nM |
|  | L-Ascorbic acid 2-phosphate (LAAP) | 50mM | 0.1mM |
|  | ITS MIX | 100X | 1x |
|  | Linoleic acid-Albumin | 100X | 1x |
| **AIM2** |  |  |  |
|  | DMEM/low glucose |  | 60% |
|  | MCDB201 Media |  | 40% |
|  | FBS (-20°C) |  | 2% |
|  | Penicillin/Streptomycin (P/S) | 100x | 1x |
|  | Insulin (rec. human) (INS) | 10mg/mL (water) | 5ug/uL |
|  | Indomethacin (IND) | 30mM (MeOH) | 50uM |
|  | IBMX | 5mM (MeOH) | 0.5uM |
|  | Dexamethasone (DXM) | 2mg/mL (5mM, EtOH) | 1uM |
|  | T3 | 10uM | 1nM |

**Table S4: Primer list**

| **Gene** | **Forward sequence** | **Reverse Sequence** |
| --- | --- | --- |
| AdipoQ | GGT CTT ATT GGT CCT AAG GG | GTA GAA GAT CTT GGT AAA GCG |
| plin1 | GCG GAA TTT GCT GCC AAC ACT C | AGA CTT CTG GGC TTG CTG GTG T |
| PPARG | AAA GAA GCC AAC ACT AAA CC | TGG TCA TTT CGT TAA AGG C |
| Oct4 | GTT GAT CCT CGG ACC TGG CTA | GGT TGC CTC TCA CTC GGT TCT |
| Pref-1 | CTT TCG GCC ACA GCA CCT AT | TGT CAT CCT CGC AGA ATC CAT |
| GATA3 | AGG GAC GTC CTG TGC GAA CT | GGT CTG GAT GCC TTC CTT CTT CAT |

**Table S5: Molecular beacon list**

| **Name** | **Sequence** |
| --- | --- |
| PPARG-799 MB | +CTT +CTC /iCy3/+CT T+CT C+GG C+CT GTG GGC CGA GAA G/3BHQ_2/ |
| PPARG-799_oligo | GGC GGA TGC CAC AGG CCG AGA AGG AGA |
| ADIPOQ-353 MB | +ACT +CCG /iCy3/+GT +TT+C A+CC GAT GTC TCC CAT CGG TGA AAC /3BHQ_2/ |
| ADIPOQ -353_oligo | CTA AGG GAG ACA TCG GTG AAA CCG GAG TAC |

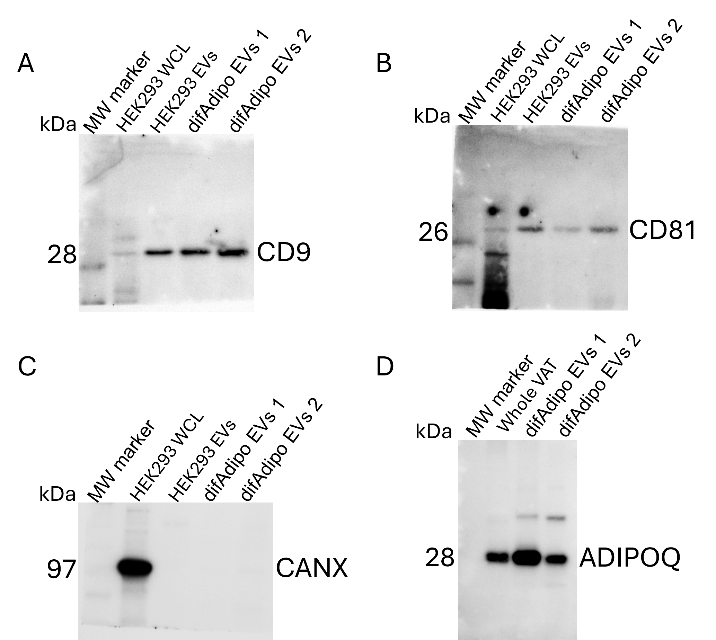

**Supplementary Figure S1.** Full-size uncropped Western blot images corresponding to the cropped blots shown in Figure 5C. EVs were isolated from differentiated adipocytes (difAdipo) of bariatric patients, referred to as EV1 (Patient 1) and EV2 (Patient 2). Each blot includes three lanes: molecular weight marker (left), difAdipo EV1, and difAdipo EV2. (A) Full-size blot for CD9 (~28 kDa), a common tetraspanin EV marker, showing positive expression in both difAdipo EV1 and difAdipo EV2. (B) Full-size blot for CD81 (~26 kDa), another EV-enriched tetraspanin, also detected in both samples. (C) Full-size blot for Calnexin (CANX; ~97 kDa), an endoplasmic reticulum protein used as a negative marker for EVs, which was absent in both difAdipo EV1 and difAdipo EV2, confirming minimal contamination by cellular debris. (D) Full-size blot for Adiponectin (ADIPOQ; ~28 kDa monomer and higher molecular weight oligomers), an adipocyte-specific protein, confirming the adipocyte origin of the difAdipo EVs. These full-size images are provided to validate the specificity, band integrity, and transparency of the immunoblotting data presented in the cropped panels of Figure 5C. Only minimal linear contrast adjustment was applied, and no other image modifications were made. WCL, whole cell lysate; VAT, visceral adipose tissue
